# ezGeno: An Automatic Model Selection Package for Genomic Data Analysis

**DOI:** 10.1101/2020.09.30.319996

**Authors:** Jun-Liang Lin, Tsung-Ting Hsieh, Yi-An Tung, Xuan-Jun Chen, Yu-Chun Hsiao, Chia-Lin Yang, Tyng-Luh Liu, Chien-Yu Chen

## Abstract

To facilitate the process of tailor-making a deep neural network for exploring the dynamics of genomic DNA, we have developed a hands-on package called ezGeno that automates the search process of various parameters and network structure. ezGeno considers three different sets of search spaces, namely, the number of filters, dilation factors, and the connectivity between different layers. ezGeno can be applied to any kind of 1D genomic input such as genomic sequences, histone modifications, DNase feature data and so on. Combinations of multiple abovementioned 1D features are also applicable. Specifically, for the task of predicting TF binding using genomic sequences as the input, ezGeno can consistently return the best performing set of parameters and network structure, as well as highlight the important segments within the original sequences. For the task of predicting tissue-specific enhancer activity using both sequence and DNase feature data as the input, ezGeno also regularly outperforms the hand-designed models. In this study, we demonstrate that ezGeno is superior in efficiency and accuracy when compared to AutoKeras, a general open-source AutoML package. The average AUC of ezGeno is also consistently higher than the result of using a one-layer DeepBind model. With the flexibility of ezGeno, we expect that this package can provide future researchers not only support of model design in their analysis of genomic studies but also more insights into the regulatory landscape.

**Availability:** The ezGeno package can be freely accessed at https://github.com/ailabstw/ezGeno.

**Contact:** Dr. Chien-Yu Chen

## 1 Introduction

We have witnessed the emergence of various high-throughput techniques and data in the biological field over the past few decades. ChIP-seq^1^ is one of the most popular experimental techniques that can track the genome-wide profiles of specific instances. Using the ChIP-seq data of transcription factors (TFs) as an example, we are interested in eventually identifying the specific binding patterns and roles of TFs in the regulatory landscape. To tackle this problem, several statistics-based tools have been developed to identify the crucial peaks among all the called peak regions. Furthermore, many pattern mining and motif discovery tools have also been utilized to detect the specific patterns in a single ChIP-seq experiment. The underlying assumption behind these approaches is that the peaks within ChIP-seq data contain at least one specific binding pattern of the given transcription factor. However, the called peaks in ChIP-seq data often contain more than one type of transcription factor binding sites, for example binding sites from co-factors and other interacting partners. These subtle differences are often hard to distinguish and, as a result, traditional motif discovery methods could fail to capture such complicated motif patterns. Therefore, in order to achieve further comprehensive studies of ChIP-seq data, an analysis tool with effective model capacity and decision-making process is much desired.

Compared with conventional machine learning techniques, deep learning, owing to its impressive performance, has exploded in popularity in the fields of data science, computer vision, natural language processing and speech recognition. In exploring Bioinformatics, many researchers have also attempted to leverage deep learning’s ability to better capture the complex, and often hidden features of training data to learn the prediction models^2–5^. DeepBind^4^ is one of the pioneers of applying deep learning to analyze transcription factor binding events within experimental data, including ChIP-seq, SELEX, and protein binding microarray data. Despite the simple structure of DeepBind, which consists of a rudimentary CNN model of fixed structure, this deep-learning based model could still achieve satisfactory results for some of the ChIP-seq data of transcription factors. We hypothesize that the adopted simple structure limits the performance of DeepBind in some of the transcription factors because the resulting neural network is not sufficient to uncover pivotal local and global information from the given genomic sequences. To better address the problem, we develop a useful package called *ezGeno* to automatically build a deep learning model tailored specifically for the given ChIP-seq data. Researchers can now utilize ezGeno to quickly obtain the best neural network architecture for the given dataset, greatly reducing the time needed for tuning the network parameters manually. At the core of ezGeno is the modified implementation of efficient neural architecture search (ENAS) that enables exploring the parameter and network structure search-space in a modest amount of time. Furthermore, not only can ezGeno find the best structure of the deep neural networks, it can also assist researchers in identifying active regions of the input sequences. In this study, we show that ezGeno is the state-of-the-art methodology that outperforms other studies in detecting patterns in the given ChIP-seq data.

## 2 Methods

We begin by illustrating the entire workflow of the ezGeno package in Figure 1, where the portion marked with a red box reflects what ezGeno package automates. The automatic procedure proceeds as follows. Starting with a positive sequence file, ezGeno first produces negative sequences, yields a well-optimized (automatic model selection) prediction model, and optionally displays a visualization heatmap on the original sequences.

**Figure 1.**
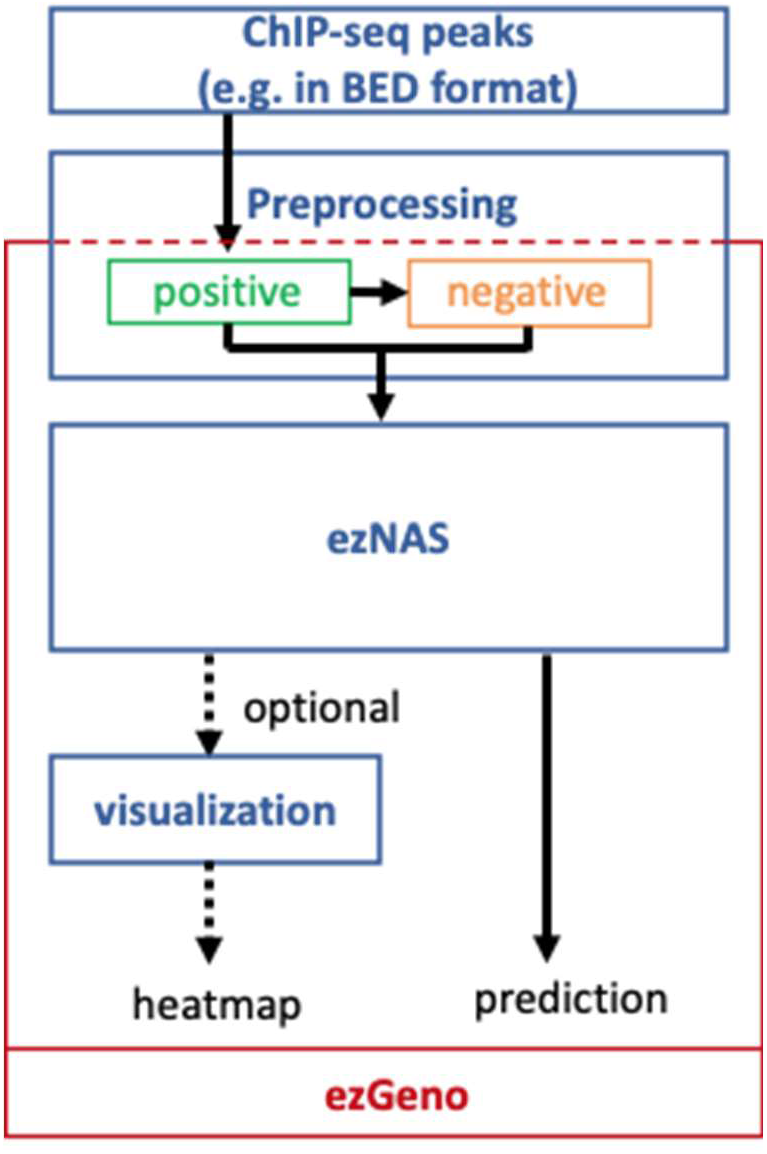
The design overview of ezGeno for genomic data analysis.

### 2.1 Data preprocessing

The data preprocessing step of ezGeno is flexible. Given a collection of positive DNA sequences or genomic features derived from ChIP-seq peaks, it can automatically generate either a corresponding set of negative instances or an augmented positive set as needed. For example, in some applications, it is needed to convert positive DNA sequences into their reverse-complement counterparts. As we will elaborate the details in the section of Applications, ezGeno deploys several handy strategies to generate negative datasets so that the underlying machine learning procedure can be appropriately carried out.

### 2.2 ENAS

#### Efficient Neural Architecture Search

With its notable progress in efficiency, Neural Architecture Search (NAS)^6^ has attracted attention as an automatic model design practice and achieved state-of-the-art results in several diverse tasks. The NAS approach can eliminate the arduous task of hand-crafted model design, an extremely time inefficient process when considering all the different combinations of hyperparameters. In a typical NAS formulation, we only need to define a search space, an objective function and a target dataset. The algorithm can then be deployed to find a good model to meet our requirements. The problem of model design therefore becomes a problem of how to efficiently search for the optimal architecture in a large search space, which usually comprises at least 10^8^ architectures. Zoph et al. ^6^ proposed the first NAS framework by training a Recurrent Neural Network (RNN) controller to learn what kind of models in the search space have higher probability to perform well on the target task. Their method first samples a set of models in the search space and trains these models by evaluating their performance. Then, Reinforcement Learning (RL) is adopted to train the controller, where the performance of the sampled models is considered as reward. By design, the controller would gradually learn what combinations and structures of model architecture are important and use this knowledge to sample the next set of models. After repeating this process several times, the model with the highest probability in the search space is determined to be the best model learned by the controller. Following this work, several NAS algorithms have been proposed with different searching strategies, different objectives, and different target datasets^7,8^. Besides RL, Evolutionary Algorithm (EA) and differentiable architecture are also frequently used in NAS^9,10^.

Efficient Neural Architecture Search (ENAS) proposed by Pham et al. ^7^ further improves NAS by sharing weights between sampled models to speed up the convergence rate of each model and by evaluating the model’s performance using mini-batches of data. Via constructing a large graph to represent the search space, sampling a model from the search space is reduced to sampling a sub-graph from the graph. The large graph can be seen as a “supernet” which contains all the possibilities of the target model, and the weights of the same structures are shared between different models. This weight-sharing strategy successfully speeds up the searching process about 1000x on image classification tasks. When developing ezGeno, we introduced the ENAS technique into our method and implemented it as a light version named ezNAS. Briefly speaking, ezNAS is different from ENAS on sampling strategy and residual connection. As we aim to analyze genome data, we consider the search space with networks composed of 1D convolution for better extraction of sequence information. Furthermore, we evaluate the model performance by AUC metric rather than accuracy to deal with data imbalance and for consistency to our final objective. A model which fits the provided dataset well can be identified, and our experiments showed that it can consistently surpass DeepBind results with better run-time efficiency. In the following subsections, we detail how ENAS is incorporated into the proposed ezGeno.

#### Search Space

Due to the sequential property of DNA sequences, 1D convolution is suitable for extracting local information. We therefore focus on the architecture of 1D Convolutional Neural Networks (CNNs) in our ezGeno framework. First, we search for the best *N*-layer CNN, which is composed of a set of possible operations and several residual connections. The possible operations can be any combination of 1D convolutional layers with different filter sizes and different dilation factors. We can define the operation set 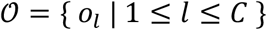 with *C* possible operations. In other words, the target network has *C* candidate operations in every layer, which generates *C*^*N*^ possible combinations. In addition, the residual connections between layers that make up the output feature map in each layer can be aggregated with one of the previous outputs, leading to *N*! possible combinations. This sums up to a total of *C*^*N*^ × *N*! models. As for tasks with multiple sources of training data, such as the second application introduced in this study (AcEnhancer), we construct a network for each data source, but perform our search jointly. The model will aggregate the features extracted from each individual source to make the final prediction. As a result, the corresponding search space includes (*C*^*N*^ × *N*!)^*K*^ models, where *K* is the number of data sources.

Any model in the search space can be encoded into a sequence of numbers. The sequence includes two types of information, the choice of candidate operation and the choice of residual connection. Since the decision of previous layers will affect the decision of the next layer, the order of model encoding is performed layer by layer, with each layer first choosing the operation, then choosing which layers to connect with. The described encoding process takes the following form:

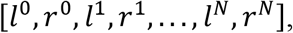

where *l* is the operation identifiers (ID) and *r* is the connected layer ID.

#### Search Process

The searching process can be divided into three stages: (1) training the supernet; (2) training the controller; and (3) deriving and evaluating the best architecture.

1. Training the supernet: We use a directed acyclic graph (DAG) to represent our search space 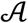 and construct a supernet denoted as 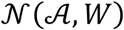 with weights *W*. We optimize *W* by the following equation:

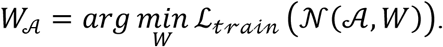 This optimization becomes inefficient in a large search space where results in a massive supernet. To deal with this problem, we optimize the weights of the supernet by architectures 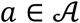 following the single path one-shot approach^10^:

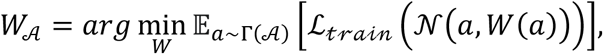

where Γ is a prior distribution. In ENAS, Γ is learned by a controller network and thus the supernet and the controller must be trained alternately. To decouple these two training processes, ezNAS only performs uniform sampling when training the supernet. We construct a sampler to sample a sequence of numbers where the encoding of the model 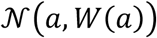 is as described previously. In every iteration, we sample a model from the supernet and train it with a batch of training data. The weights of the supernet can then be updated correspondingly.
2. Training the controller: In the ezGeno framework, the architecture search objective can be written as follows:

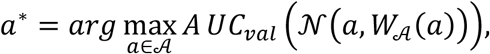

where 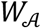 is trained in the previous stage and is fixed in this stage. A validation dataset different from training dataset is used for generalizability. However, the objective is still hard to optimize directly, especially in a large search space. We use a controller to help retrieve the best model. Due to the dependency between layers, we construct the controller using LSTM cells^11^. LSTM is designed to handle time series data and is suitable to learn sequential decisions. To train the controller, we rewrite the objective:

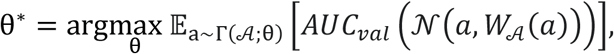

where θ is the weights of the controller. In every time step, we decode the hidden state to get the probability of each choice and make a decision by sampling from the probability. After several time steps, a model can be decided, and we evaluate its performance with a batch of validation data. We then take the performance as rewards and compute the gradient of the controller by REINFORCE^12^. The controller can be updated according to the gradients and the optimizer.
3. Deriving and evaluating the best architecture: After the training of the supernet and the controller, we repeat the sampling process but only keep the model with the highest probability of being the best model:

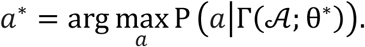 Having decided the *optimal* 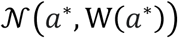, we perform training from scratch to ensure the constructed network could achieve its best performance to the target task.

### 2.3 Visualization

In ezGeno, we adopt a gradient-based localization method, Gradient-weighted Class Activation Mapping (Grad-CAM)^13^, to show which regions of the original sequences are important for performing predictions and then can be considered when constructing possible motifs. Since a deeper convolutional layer in the model extracts more high-level semantics while keeping important spatial information, we can locate the region in the input data which may affect the predictions through the feature map activations in the final convolution layer. Considering a convolutional neural network with several convolutional layers, global average pooling layer and fully connected layer, *A*^*k*^ is the feature map activations. For a given class *c*, *y*^*c*^ is the predicted score of the class and is computed by:

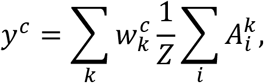

where *w* is the class feature weights, *i* is the index over width dimension and *k* is the index over *K* feature maps. We can then compute the importance weight 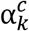 by

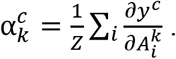

Finally, the Grad-CAM can be obtained by

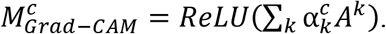

Through the Grad-CAM interpretation, we can further analyze our predictions by visualizing our model’s decision-making process.

## 3 Target Applications

To demonstrate the usefulness of ezGeno in genetics analysis, we have considered two important applications and applied ezGeno on these applications by properly defining the search space of network structure. The efficiency of ezGeno is further justified by comparing with AutoKeras in terms of performance and speed. We explain the details of how to carry out the experiments and the comparisons in the next two sections.

### 3.1 TF Binding (Task name: TFBind)

Transcription factors (TFs) are regulatory elements that can alter gene expression profiles by binding to specific regions of the genome. In order to discover the specific binding patterns of different transcription factors, several functional assays have been developed in the past few decades. ChIP-seq is one of the most widely used experimental methods to identify the binding events for a transcription factor under the given genomic context. Recently, a deep learning framework, coined as DeepBind, was proposed to demonstrate the possibility of identifying the binding motif of the given ChIP-seq data for a transcription factor. The performance of DeepBind on different transcription factors varies because the intrinsic properties of transcription factors are non-identical.

#### Dataset

We utilized the TF ChIP-seq data to examine the performance of ezGeno. We first downloaded the data of 20 transcription factors (Table 1 and Table 2) from the curated database of DeepBind^4^ and split the ChIP-seq called peaks from the ENCODE database^14^ into three categories: A, B and C. Category A comprises the top 500 even-numbered peaks while Category B incudes the top 500 odd-numbered peaks. The rest of the called peaks were collected into category C. We used category A and C as our training data and category B as the testing positive dataset. To ensure the models can be trained with enough data points, we applied random sampling with replacement to the positive data until every dataset has at least 10,000 data points. Next, we generated an equal amount of negative data based on the user’s requirements. One strategy is to generate negative instances that match the dinucleotide composition of the positive data.

**Table 1.** The top-10 worst performing cases in DeepBind.

| TFs | Cell type | URLs |
| --- | --- | --- |
| RBBP5 | H1 | https://www.encodeproject.org/experiments/ENCSR000AQC/ |
| RXRA | Gm12878 | https://www.encodeproject.org/experiments/ENCSR000BJD/ |
| FAM48A | HeLa-S3 | https://www.encodeproject.org/experiments/ENCSR000ECQ/ |
| WRNIP1 | GM12878 | https://www.encodeproject.org/files/ENCFF679UWZ |
| SAP30 | K562 | https://www.encodeproject.org/files/ENCFF235UDJ |
| BDP1 | K562 | https://www.encodeproject.org/files/ENCFF002CVP |
| CHD1 | GM12878 | https://www.encodeproject.org/files/ENCFF151MEY |
| HDAC1 | K562 | https://www.encodeproject.org/files/ENCFF016MVN |
| HDAC6 | K562 | https://www.encodeproject.org/files/ENCFF669QKC |
| SUZ12 | H1 | https://www.encodeproject.org/files/ENCFF389SPE |

**Table 2.** The top-10 best performing cases in DeepBind.

| TFs | Cell type | URLs |
| --- | --- | --- |
| BATF | GM12878 | http://tools.genes.toronto.edu/deepbind/D00741.001/index.html |
| CTCF | GM12801 | http://tools.genes.toronto.edu/deepbind/D00328.018/index.html |
| HNF4A | HepG2 | http://tools.genes.toronto.edu/deepbind/D00446.009/index.html |
| JUND | HepG2 | http://tools.genes.toronto.edu/deepbind/D00776.005/index.html |
| MAFF | HepG2 | http://tools.genes.toronto.edu/deepbind/D00501.004/index.html |
| NFE2 | K562 | http://tools.genes.toronto.edu/deepbind/D00535.004/index.html |
| NFYB | K562 | http://tools.genes.toronto.edu/deepbind/D00789.003/index.html |
| NRF1 | HeLa-S3 | http://tools.genes.toronto.edu/deepbind/D00559.006/index.html |
| POU5F1 | H1-hESC | http://tools.genes.toronto.edu/deepbind/D00795.001/index.html |
| RAD21 | A549 | http://tools.genes.toronto.edu/deepbind/D00796.001/index.html |

#### Architecture

We depict our network in Figure 2, where the super-network design of our TF binding model is illustrated in Figure 2(a). At each summation node, the network decides whether or not the current output should be aggregated with that of a previous layer, a concept taken from ResNet. These probabilistic decisions are denoted using the dotted lines and can be chosen at most once, or excluded at all. Solid lines represent the original flow of the network that the model must follow. The paired numbers *f* × *d* in green indicate there are *f* feature maps, each of which has dimension *d*. Note that the aggregation does not cause any change in dimensionality since it is simply a direct summation of equal-length vectors. Figure 2(b) shows the components of each convolutional unit and the possible hyperparameters of the unit to be decided. As an example from Figure 2(b), the convolution unit is specified from totally 10 combinations of the two types of parameters. During training, the controller decides what parameters each convolution unit will choose. In the TF binding task, every convolution unit has a total of 10 design choices, in each of the three layers, with every output having the option to connect with the outputs of previous layers. This yields a total search space of 10^3^ × 3! = 6000 models.

**Figure 2.**
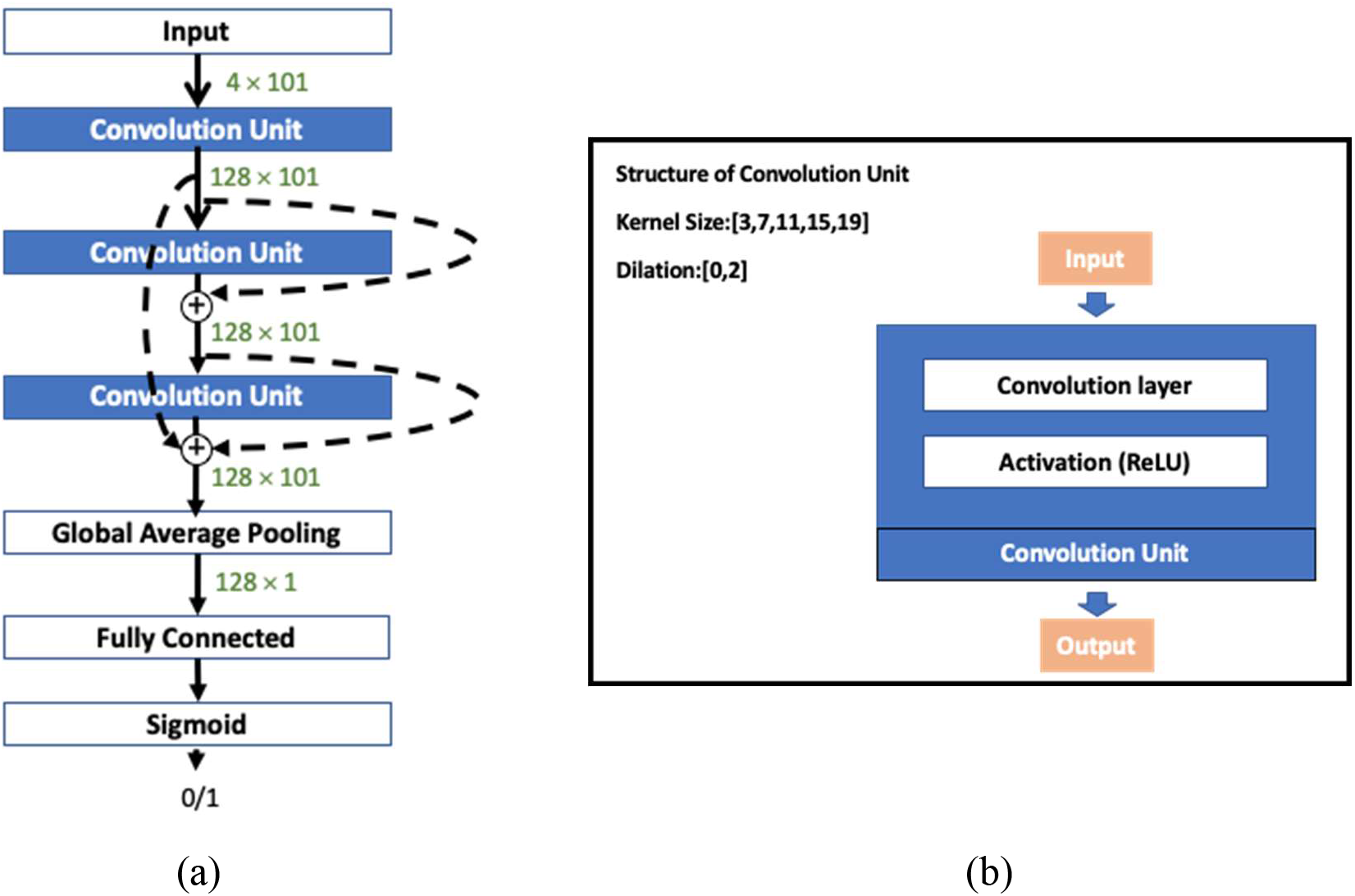
The search space of the task: TF binding (TFBind)

#### Training process

The supernet is constructed first by performing weight sharing training, and then by using reinforcement learning to learn specific model performances in order to sample the best performing model. During the search process, the controller aims to decide the appropriate convolution filter size and the aggregation to the output of a previous layer. Table 3 lists the default hyperparameters used for the TF-binding task in ezGeno.

**Table 3.** The hyperparameters of TFBind and their default values.

| Hyperparameters of the task TFBind | Default |
| --- | --- |
| task | TFBind |
| Batch size | 64 |
| supernet learning rate | 0.01 |
| Supernet epochs | 100 |
| optimizer | sgd |
| controller learning rate | 0.001 |
| controller_optimizer | adam |
| cstep | 2000 |
| learning rate | 0.01 |
| momentum | 0.9 |
| weight decay | 5e-4 |
| epochs | 100 |
| layers | 3 |
| feature_dim | 128 |
| conv_filter_size_list | [3,7,11,15,19] |

#### Visualization

In previous work, Alipanahi et al. used convolution filters as motif detectors to uncover motifs, but some cases could not be predicted satisfactorily^4^. Using convolution filters as motif detectors might limit the discovery of complicated patterns. Lanchantin et al. explored a saliency map to detect motifs, but their method turned out to be highly sensitive to noise^15^. In ezGeno, we instead consider Grad-CAM^13^ to learn the positional importance within each sequence. The Grad-CAM operation yields a gradient vector whose score represents the importance of each input sequence. We then use a sliding window and pass it to a pooling layer from the gradient vector. If the score is greater than the threshold, we will peak these sub-sequences, while extending the sub-sequence if the other position in the sliding window satisfies the requirement. We also collect the sub-sequences whose scores surpasses the threshold after pooling and save them in FASTA format. This file can be treated as the input to a motif discovery tool^16^ to generate motifs in the sub sequences.

### 3.2 Active Enhancer (Task name: AcEnhancer)

In addition to transcription factors participating in transcription, regulatory elements in non-coding regions also play essential roles in regulating the gene expression such as promoters, enhancers, and so on. But the mechanism for enhancers is still an open question for now. Previous studies showed that many variants in non-coding regions also have been associated with various diseases during genome-wide association studies. If we understand enhancer function more, we can know more about the causes of diseases. In the following, we demonstrate ezGeno can be applied to many aspects in genetics including enhancer prediction tasks.

#### Dataset

Many studies showed that certain histone modifications such as H3k27ac have been associated with increasing the likelihood of activating transcription and therefore defined as an active enhancer mark^17–19^. In this task, we chose H3k27ac as active enhancers and collected H3k27ac ChIP-seq data of four cell types as a positive dataset from the ENCODE database^14^. In addition to the genomic sequences, DNase feature data may also help to predict enhancers. We then extracted DNase feature data overlapped with the upstream and downstream within 2500 bps of the selected training instance as the input feature. We set 200 bps in one bin and divided these 5000 bps into 25 bins. Therefore, the DNase signal vector for a single training instance will be 25×1. As for the prediction target, enhancer activity, the h3k27ac ChIP-seq data was used to label the instance. As for the negative data, we randomly selected the negative instance outside the positive-defined regions in the genome while maintaining a positive to negative ratio of 1:10.

#### Architecture

For the active enhancer task, we have two types of data as the input features: the sequence data and the DNase data. We constructed a two-branch model, each branch processing one type of data. After forwarding to the final convolution layer of each branch, the features will be concatenated and forwarded into fully connected layers to make the final predictions. Figure 3 illustrates the architecture overview of our AcEnhancer model. The structure of the convolution unit in the DNase branch has 6 operation choices, while the structure of the convolution unit in the sequence branch has 10 choices. The design of the convolution unit is similar that shown in Figure 2(b). After constructing the supernet, ezNAS can then be applied to both the two branches simultaneously. Table 4 lists the default hyperparameters used in this task.

**Figure 3.**
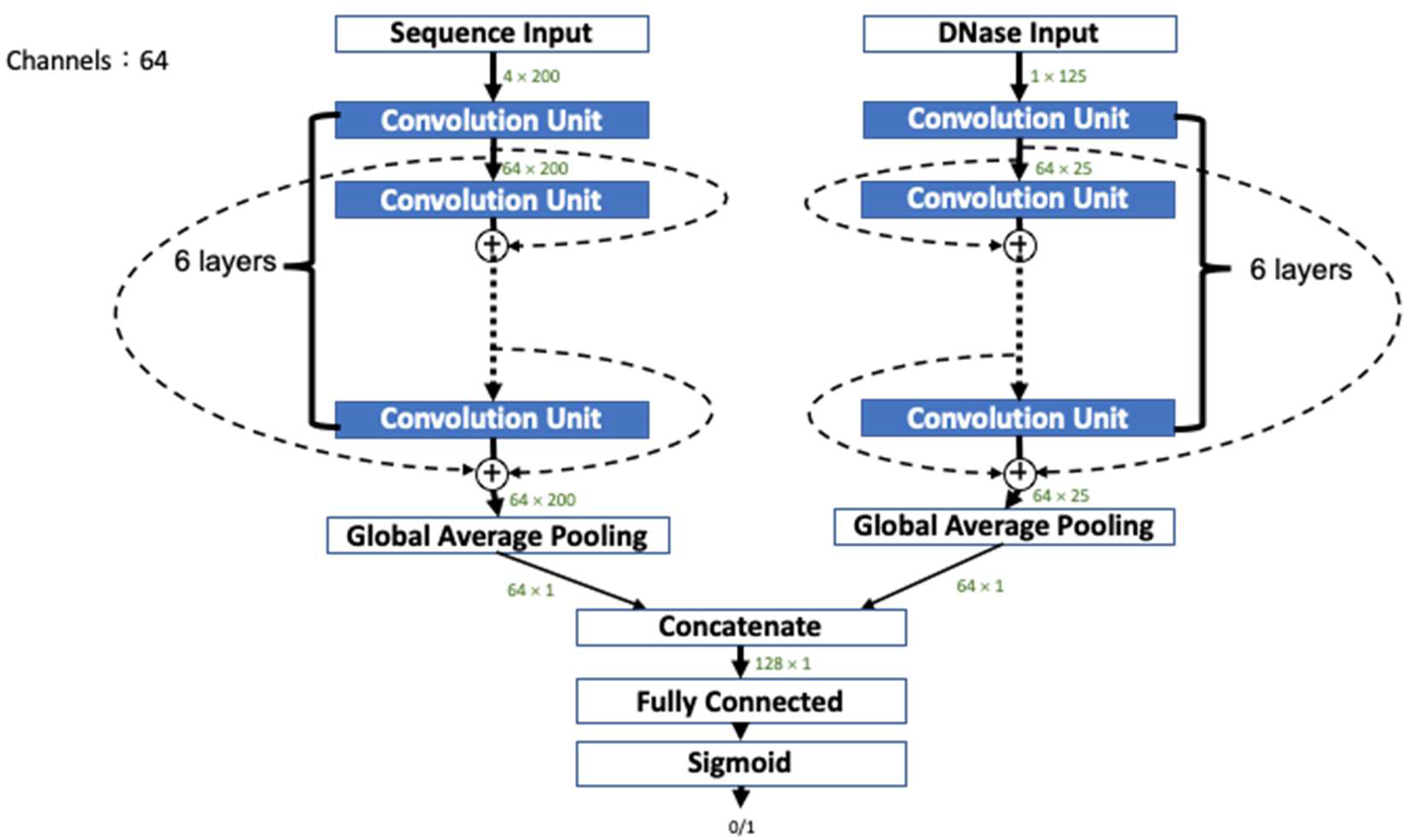
The search space of the task: predicting enhancer activity (AcEnhancer).

**Table 4.** The hyperparameters of AcEnhancer and their default values.

| Hyperparameters of AcEnhancer | Default |
| --- | --- |
| task | AcEnhancer |
| Batch size | 128 |
| supernet learning rate | 0.01 |
| Supernet epochs | 100 |
| optimizer | sgd |
| controller learning rate | 0.001 |
| controller_optimizer | adam |
| cstep | 2000 |
| learning rate | 0.01 |
| momentum | 0.9 |
| weight decay | 5e-4 |
| epochs | 100 |
| layers | 6 |
| feature_dim | 64 |
| conv_filter_size_list | [3,7,11,15,19] |
| dNase layers | 6 |
| dNase_feature_dim | 64 |
| dNase_conv_filter_size_list | [3,7,11] |

## 4 Results

In this section, we demonstrated the effectiveness of the ezGeno framework. In order to ensure the performance of ezGeno, we compared ezGeno with other existing tools. We found that ezGeno performed better than existing tools such as AutoKeras in the two tasks described previously. Moreover, the efficiency of ezGeno for searching model architectures is much more superior than AutoKeras. Details are listed in the following paragraphs.

We compared ezGeno with AutoKeras^20^, which is a popular open source framework for NAS. AutoKeras utilizes Bayesian optimization to guide the network morphism for efficient neural architecture search. It provides a set of APIs for different types of applications to quickly adopt AutoKeras for NAS. For genomic data analysis, we treated genomics data as images and utilized the *ImageClassifier* API. The trial number was set to 100 and 4 for the TF Binding and Active Enhancer task, respectively. We only set the trial number to 4 for the Active Enhancer task because of the long searching time on a large dataset. For each architecture trial, we used the early stopping criterion with the maximal threshold of 10. Data was preprocessed in the same way as ezGeno. For the TF Binding task, we encoded each ChIP-seq data into one-hot vectors. For the Active Enhancer task, we encoded the sequence data same as the TF Binding task and concatenated the DNase data to the one-hot vectors of sequence data directly.

### 4.1 TF Binding (Task name: TFBind)

From the dataset provided by DeepBind, we selected the top-10 worst performing cases in DeepBind to evaluate our experiments (Table 1). We used these 10 cases to determine which methods provided the best results. All the experiments were repeated 10 times and the average and standard deviation of the results were reported. In most of the cases, we could finish the whole searching and training process within 10 minutes on a Titan-RTX GPU, which is much faster than using AutoKeras.

Figure 4 compares the AUC of 3 different frameworks: ezGeno, AutoKeras, and DeepBind. As shown, the ezGeno models outperform AutoKeras and DeepBind models by a significant portion in all selected cases. The performance can reach an AUC of 0.8 in the experiments on SUZ12, SAP30, RXRA and RBBP5. Most of the others also have better results which are not less than 0.7. To be more specific, the models searched by ezGeno can improve AUC metric up to 0.17, compared with the DeepBind results.

**Figure 4.**
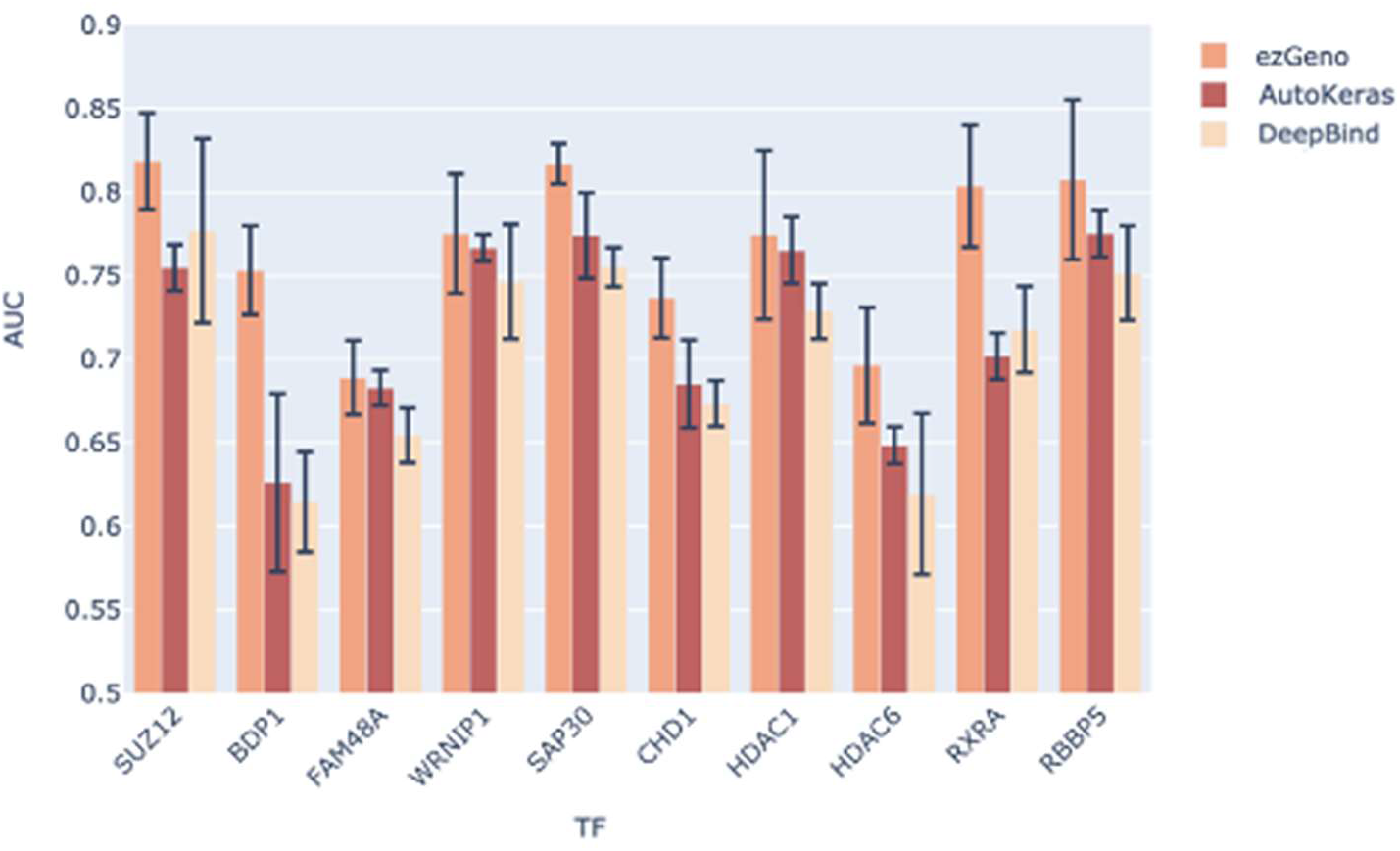
The performance of ezGeno on 10 difficult TFs, in comparison with AutoKeras and one-layer DeepBind.

Figure 5 compares the training time for Neural Architecture Search. We can see that ezGeno far outperforms AutoKeras, with an average 5 times faster than AutoKeras. ezGeno can even speed up 20 times in an extreme case.

**Figure 5.**
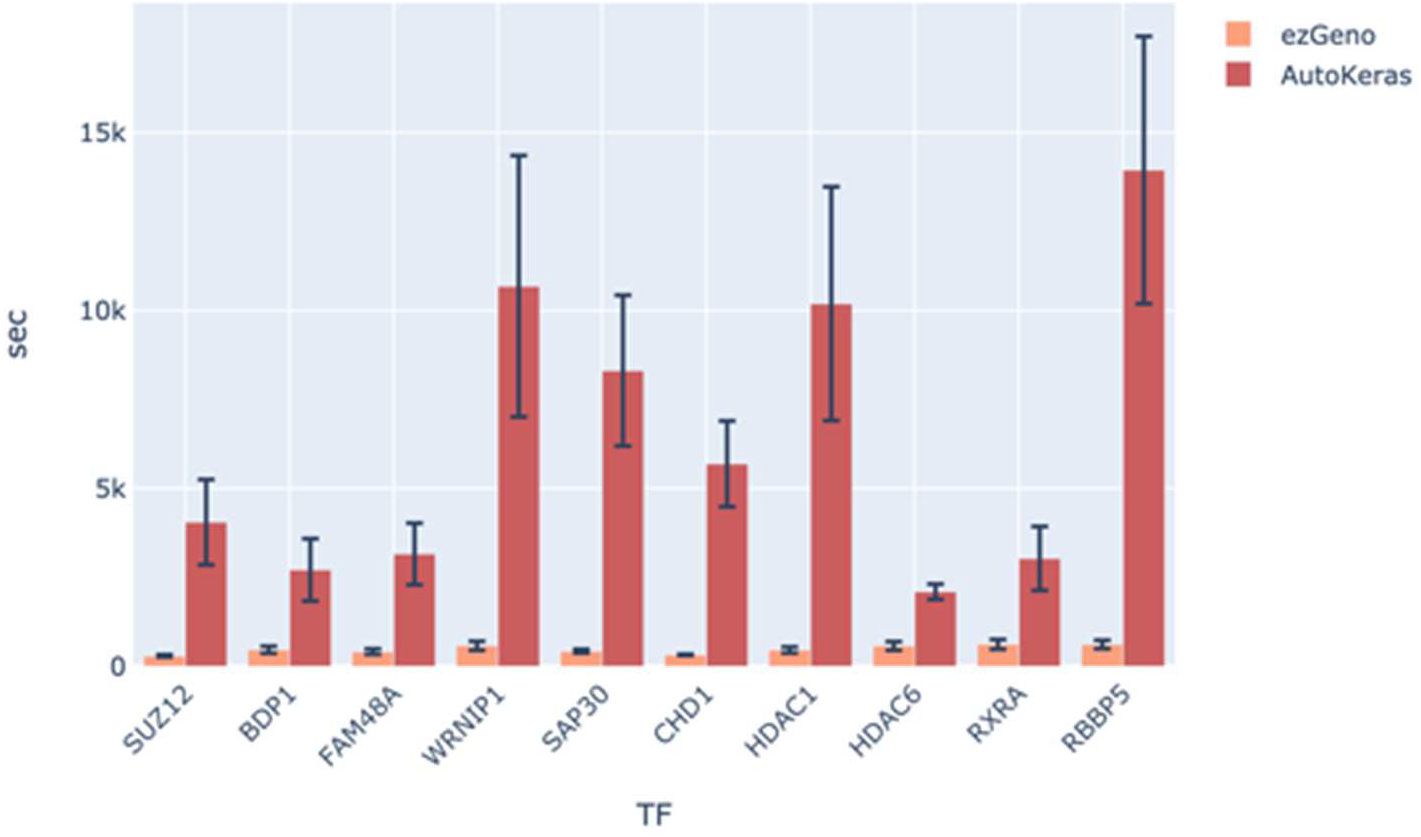
The running time of ezGeno, in comparison with AutoKeras.

We also experimented on 10 TFs (Table 2) which perform well in DeepBind (over 0.9 AUC) as a sanity check to see if ezGeno can reach the same level of performance. The results are shown in Figure 6. We noticed that the overall performance of ezGeno is similar to DeepBind and even surpasses DeepBind in some cases such as HNF4A and POU5F1.

**Figure 6.**
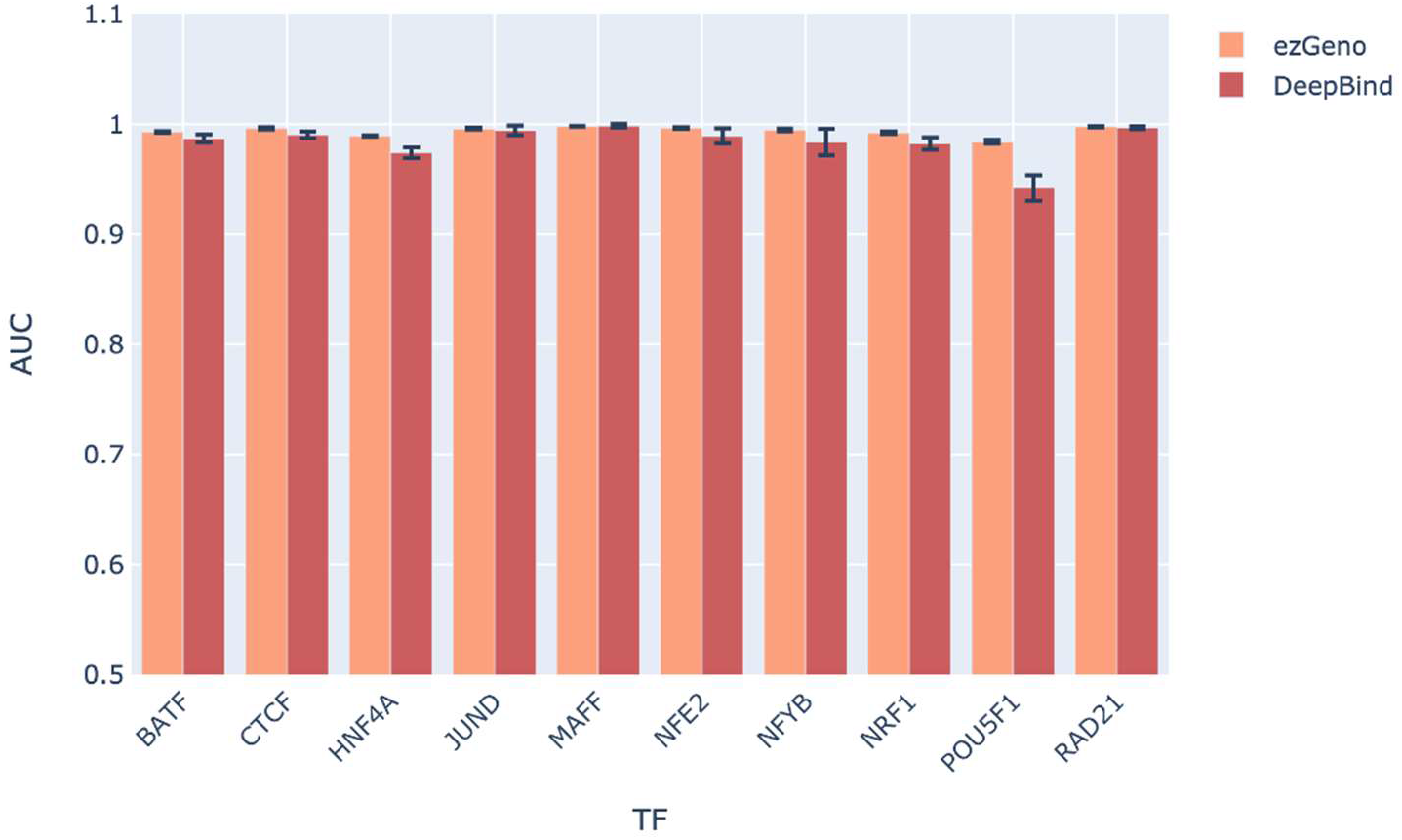
The performance of ezGeno on 10 easy TFs, in comparison to one-layer DeepBind.

In order to further understand how our models make predictions, we used Grad-CAM to visualize hotspots within the sequences. These hotspots can correspond to the segments that the model deems as high probability to be bound. Figure 7 shows the Grad-CAM of 100 positive sequences generated by the model from the MAFF TFBind experiment. We can observe that the distribution of the hotspots is concentrated arround the center of the sequences. This result is consistent with previous biological experiments.

**Figure 7.**
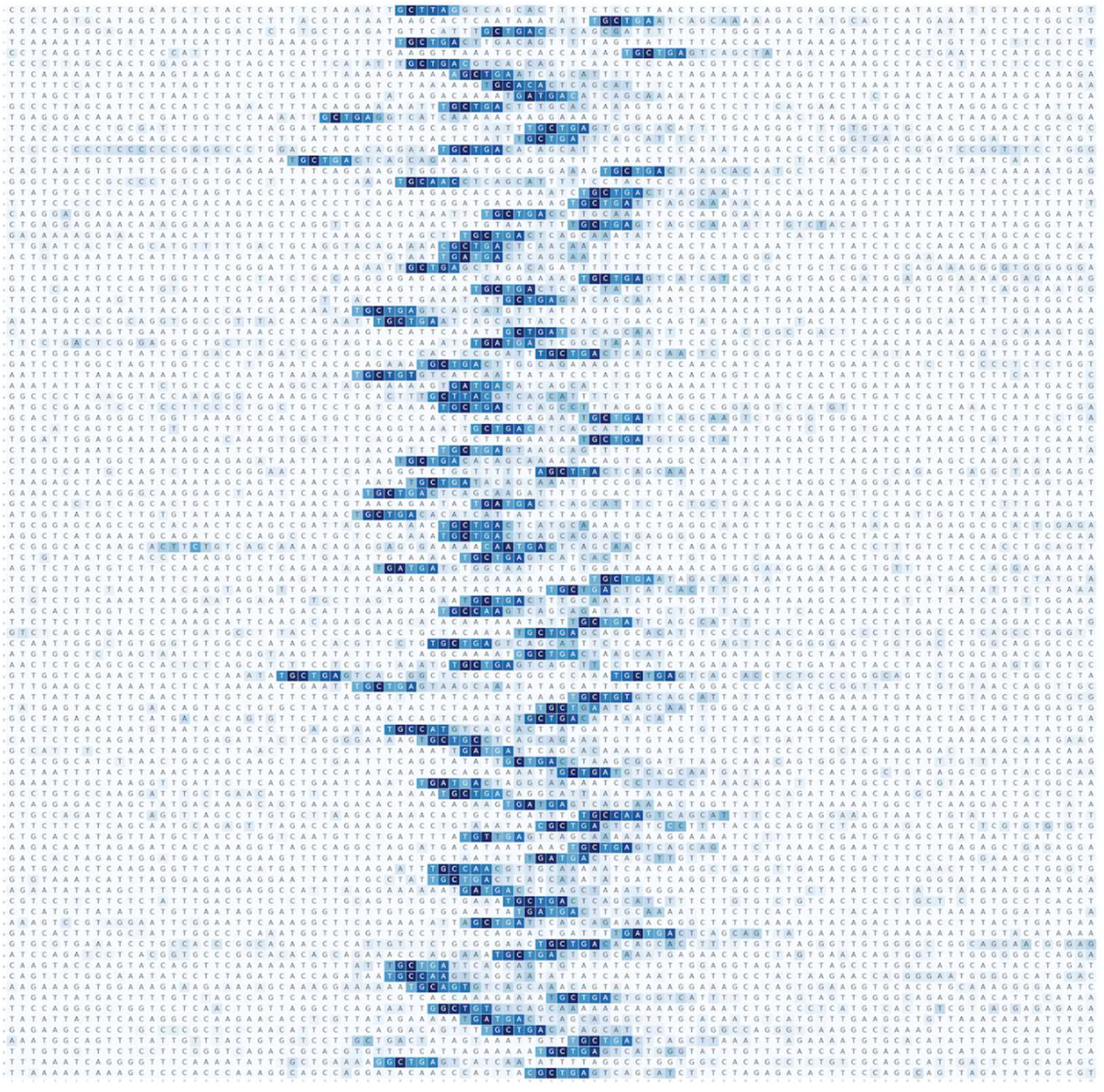
The visualization plot. The y axis represents each individual sequence, while the x axis represents position. For each sequence, the darker shade of blue, the more importance of the positions.

Through Figure 7, we can identify the important positions that our model focuses on. We set a sliding window (window size = 9) and let every sub-sequence truncated by the window pass through a pooling layer to generate a score. We then collected the sub-sequences whose scores surpass a threshold (threshold value = 0.25) and report them as FASTA format. This FASTA file can then be fed as the input of a motif discovery tool (e.g. MEME) to generate motifs. The results are shown in Figure 8. The first three columns denote the TF name, the number of sub-sequences, and the total number of site counts found by MEME, respectively. As for the last two columns, the sequence logo in the left column is the top-1 motif hit discovered by MEME from the high score sub-sequences, and the sequence logo in the right column is from the HOCOMOCO database^21^, representing the known motifs of the TFs. Comparison of the two results shows a consistency between ezGeno’s results and annotated motifs. This again indicates that we can retrieve a reliable and meaningful result using ezGeno without manual design on network structure.

**Figure 8.**
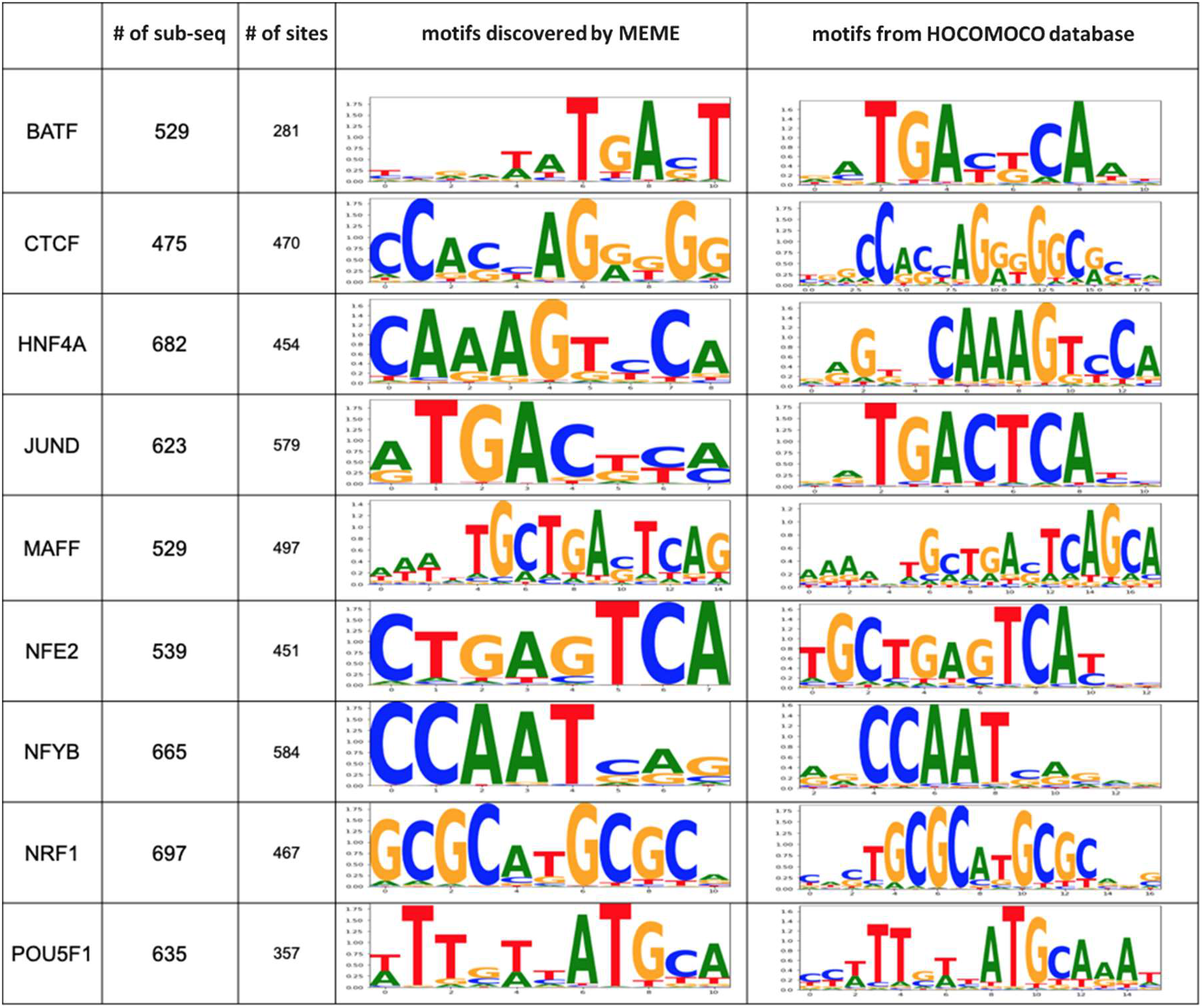
The motif discovery results on the highlighted segments retrieved by ezGeno (‘# of sub-seq’ is the number of sub-sequences reported by ezGeno through Grad-CAM; ‘# of sites’ is the number of sites discovered by MEME on the sub-sequences;).

### 4.2 Active Enhancer (Task name: AcEnhancer)

We run the Enhancer tasks using Tesla P100 GPU. The performance on the four different cell type datasets is the best when compared with other methods. As shown in Figure 9, the testing AUC of the models searched by ezGeno all surpass the hand-craft models called accuEnhancer and the searched models of AutoKeras. The comparison of search cost is shown in Figure 10, ezGeno can finish the searching process with less time consumption and is at least 3 times faster than AutoKeras. Overall, ezGeno provides more reliable and better results for the task of Active Enhancer in a much more efficient way.

**Figure 9.**
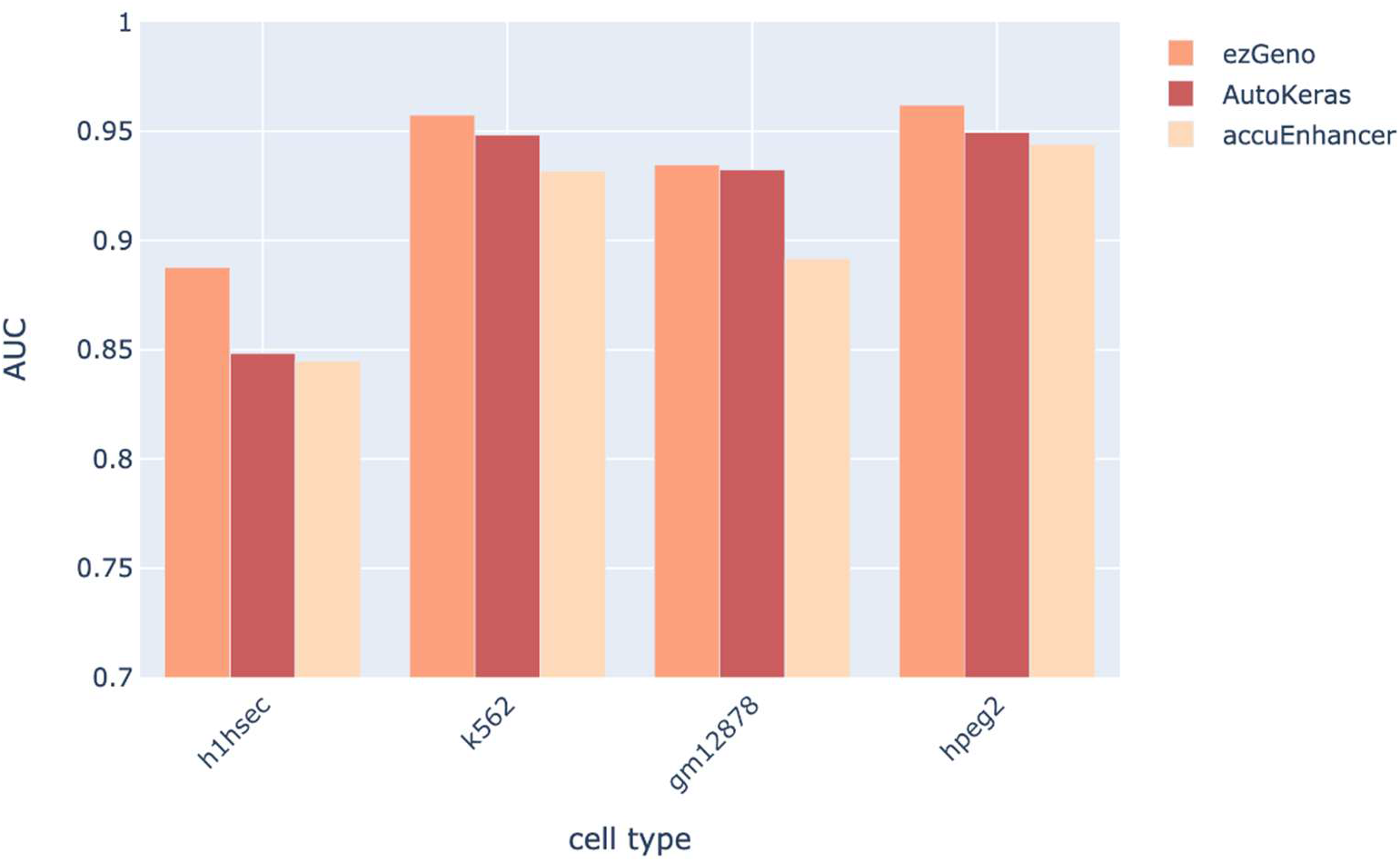
The performance of ezGeno on different cell types, in comparison with AutoKeras and the hand-craft models called accuEnhancer.

**Figure 10.**
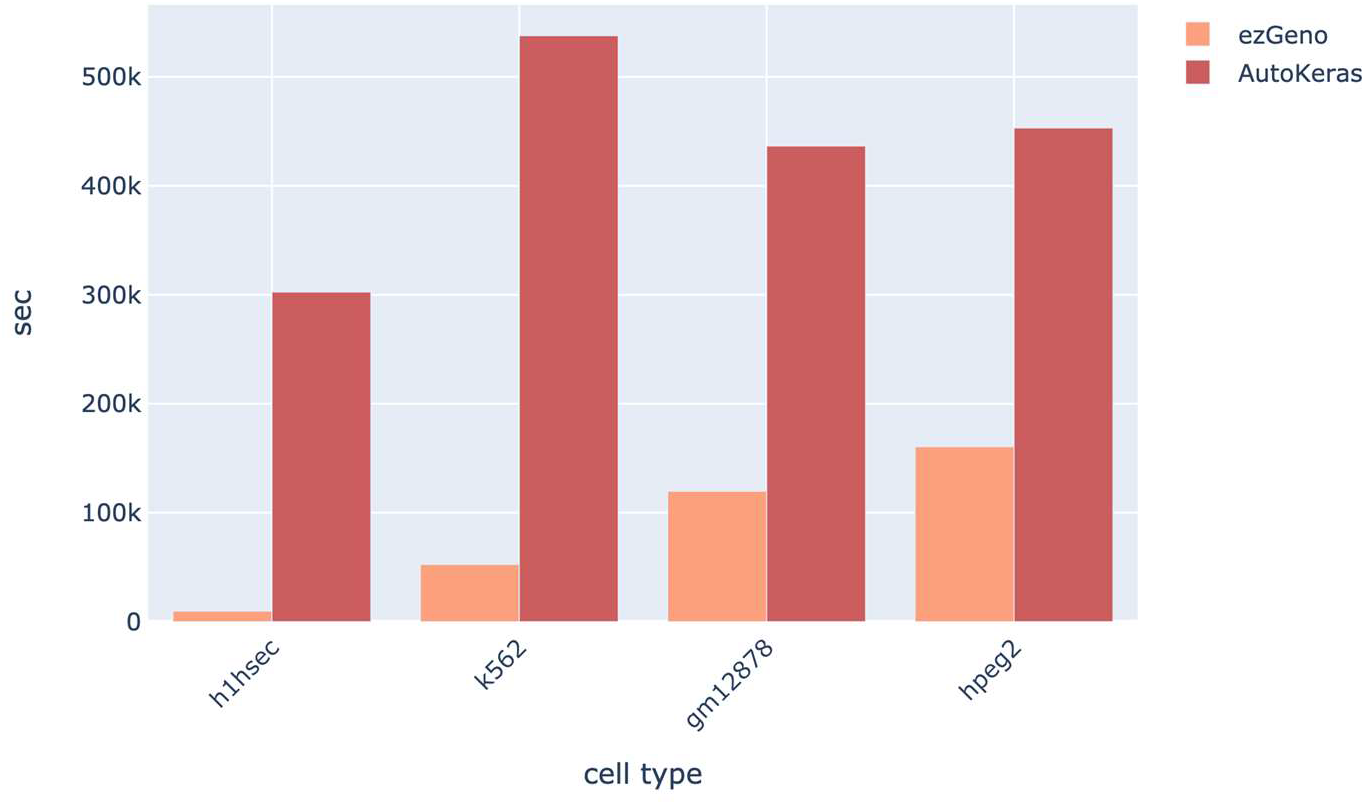
The searching cost of ezGeno in comparison with AutoKeras.

## 5 Discussions

To discuss whether the performance would benefit from a larger model, we modified the search space of TFBind to search for deeper or wider networks. In the deeper-network experiments, we increased the number of layers of the target model from 3 layers to 6 layers and reconducted the TF Binding experiments. The search space was relatively expanded into 10^6^×6! ~ 7.2×10^8^ models. Figure 11 shows the performance comparison between the 3-layer setting and the 6-layer setting. We can see that in most of the TF Binding experiments, there is no significant increase in AUC. In some of the cases such as HDAC6 and RBBP5, the best performance in the 6-layer setting between all trials cannot surpass the best one in the 3-layer setting. It suggests that there is just a little or no performance gain if the model becomes deeper. Moreover, searching with the 6-layer setting is more time-consuming. Figure 12 shows the running time of the 3-layer and the 6-layer setting. The running time increases significantly under 6-layer setting in all cases. The reason is that with the same training epochs, larger models need more time to train due to more parameters. Considering both the efficiency and the performance, we decided to make 3-layer as our default hyperparameter for the application of TFBind.

**Figure 11.**
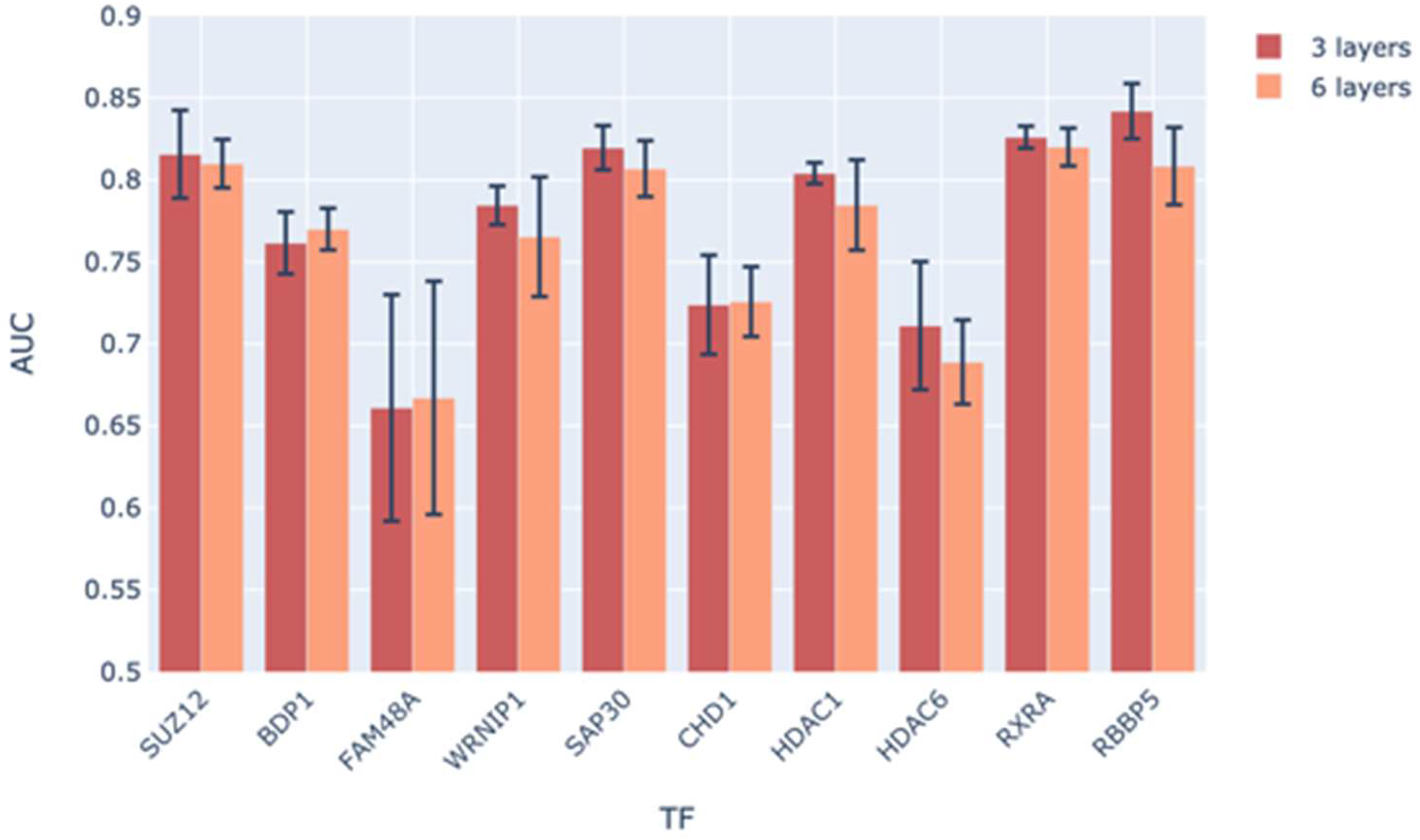
Comparison of a 3-layer ezGeno versus 6-layer ezGeno in the task: TFBind.

**Figure 12.**
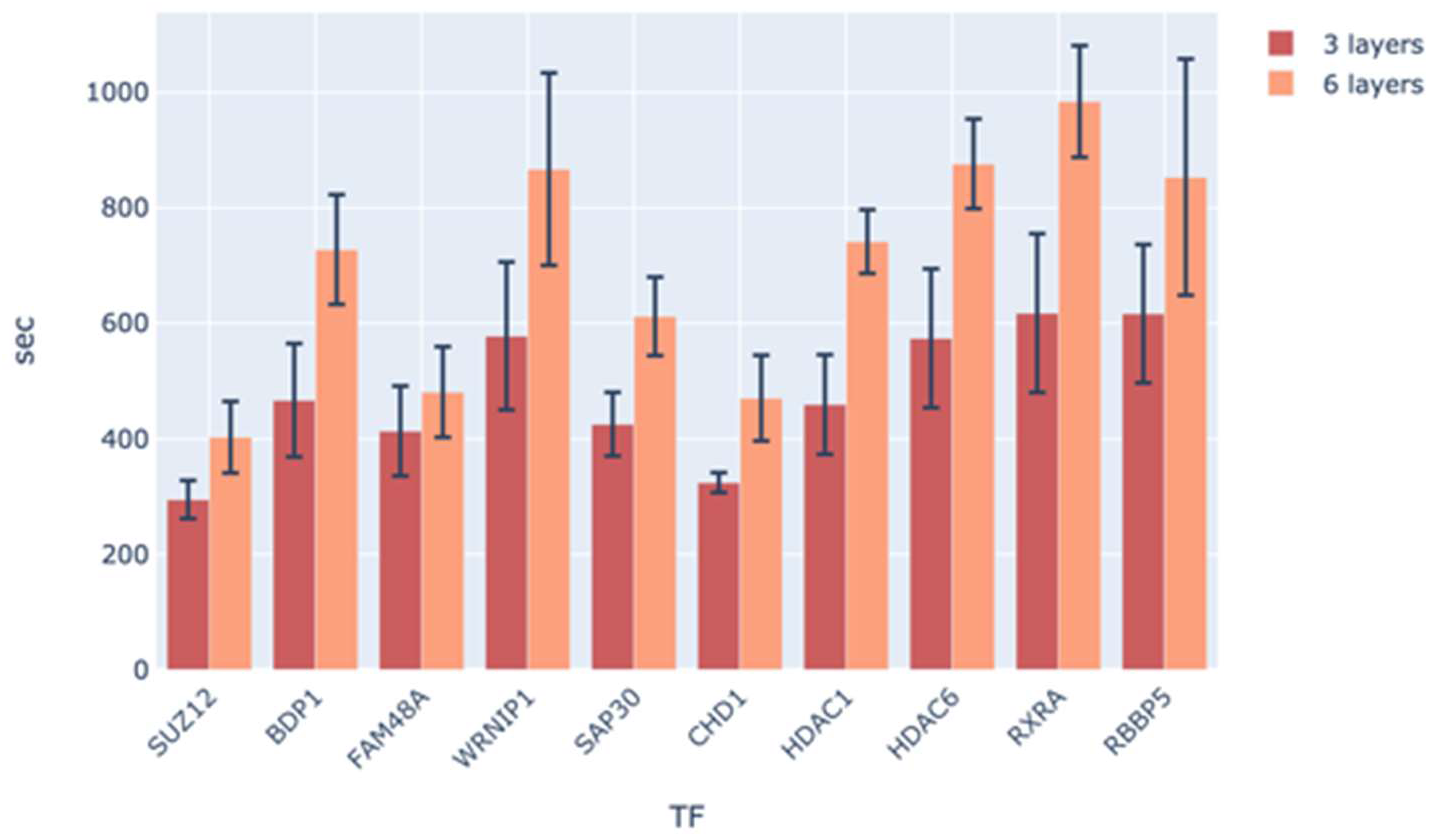
The running time of ezGeno in different settings: 3-layer versus 6-layer.

As for the wider-model test, we modified the number of channels in each layer. We constructed models with 16, 64 and 128 channels, respectively, to see the effect of channel increase. Figure 13 shows the results. In most of the cases, the testing AUC becomes higher with the number of channels increasing. As for the training time shown in Figure 14, due to the early stopping mechanism, the training time is not proportional to the model size. We can see that most of the models with 128 channels converge faster and thus need less time to achieve satisfactory results. The observation leads us to prefer the model wider rather than deeper to boost the performance.

**Figure 13.**
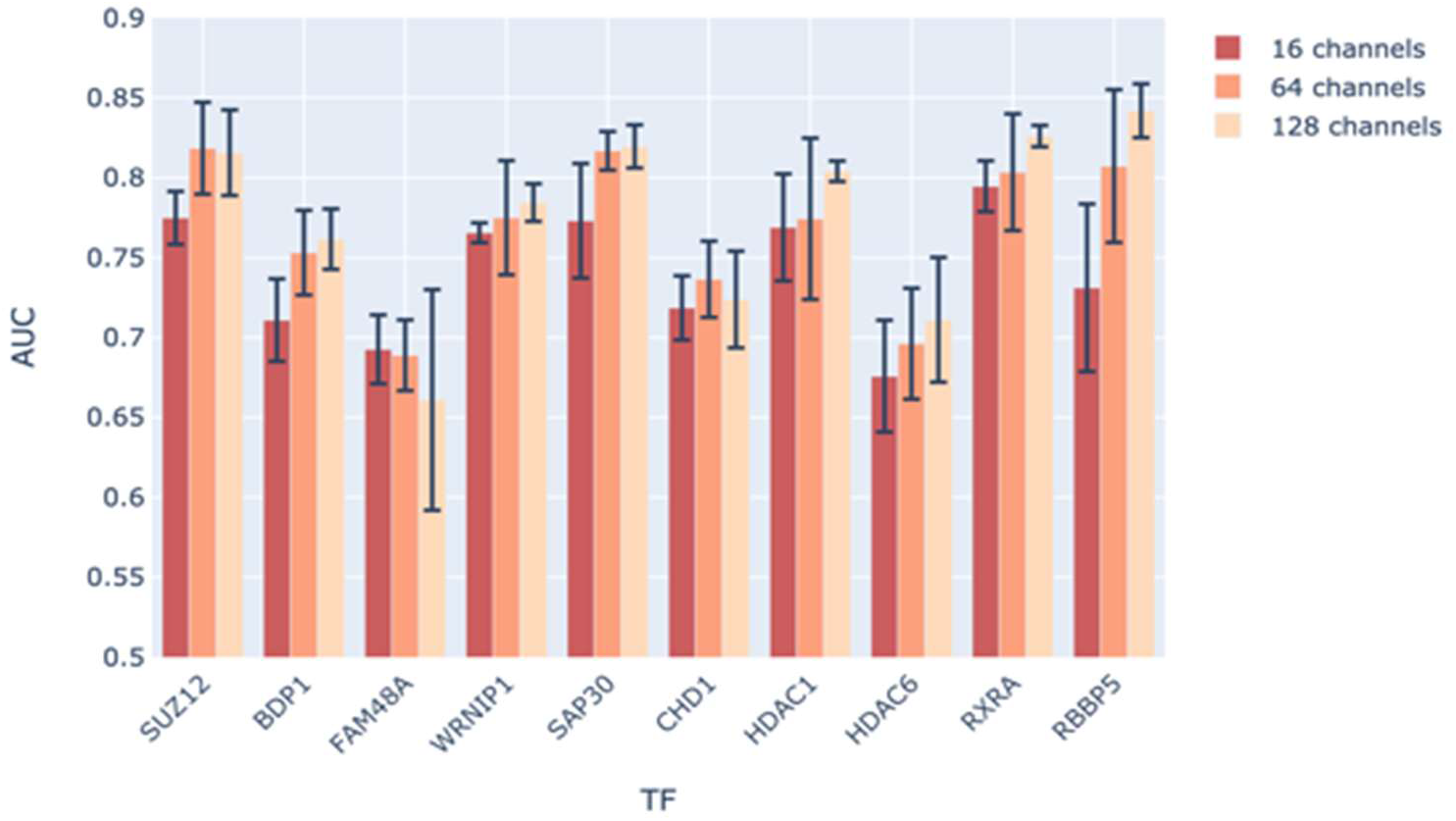
Comparison the performance of ezGeno with different numbers of channels.

**Figure 14.**
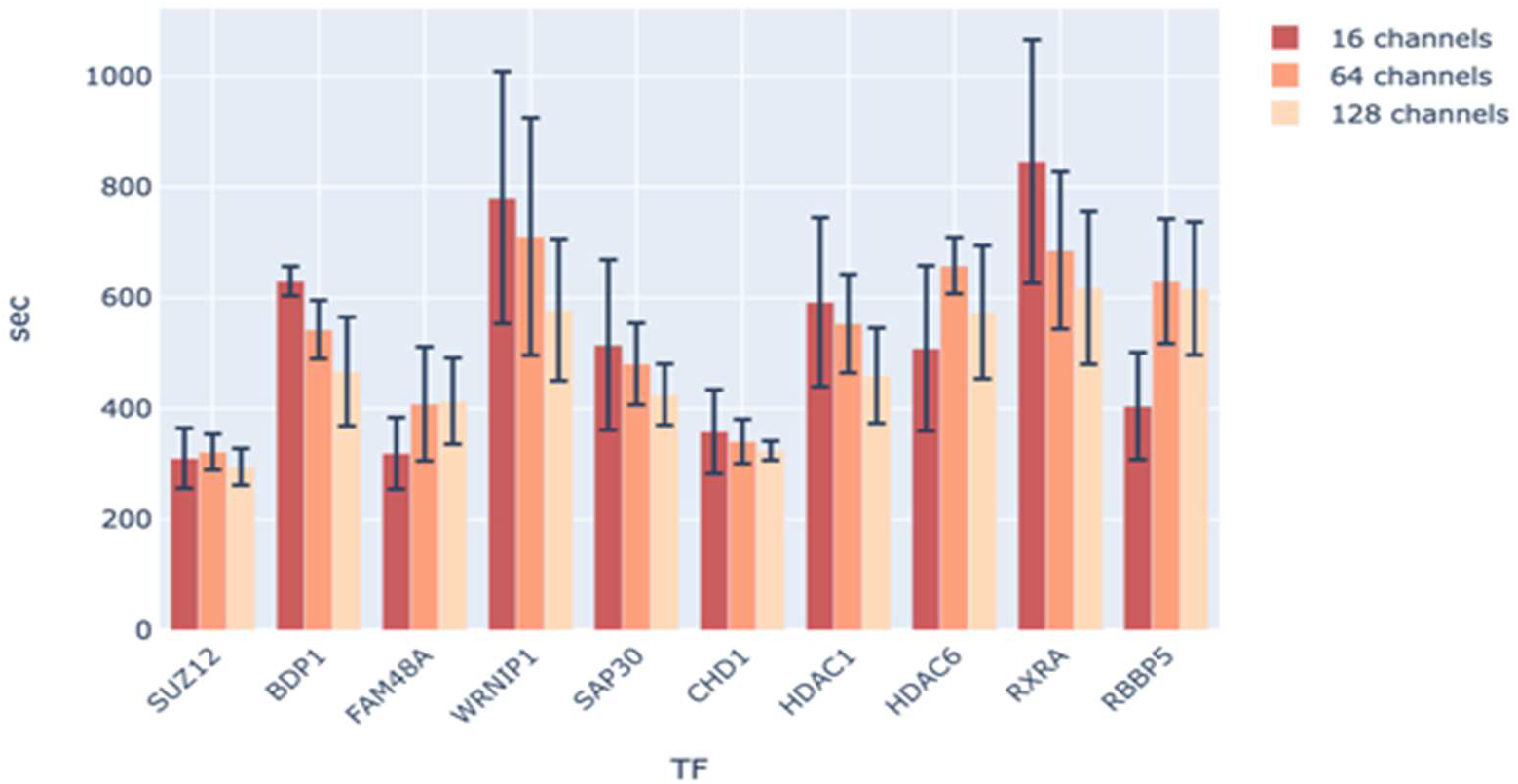
The running time of ezGeno in different numbers of channels.

## 6 Conclusion

In this work, we proposed an automatic framework, ezGeno, to help users run genome-related analysis on their own data without heavy workload. To be more specific, ezGeno can automatically perform data preprocessing, model searching, prediction, and model interpretation. After data preprocessing, users can obtain negative datasets based on positive datasets. Furthermore, ezGeno exhibits a high degree of flexibility that it can process not only genomic sequence data but also any kind of 1D genomics such as histone modifications, DNase feature data, or even combinations of these data. During training, ezGeno exploits the NAS algorithm to help users find high performance models in different tasks. Our experiments showed that ezGeno can provide reliable results which outperform previous works with less search cost. Finally, ezGeno can generate heatmaps by the Grad-CAM algorithm to help users explain their findings. These results showed a high consistency with the biological experiments. In conclusion, the promising results shown in this study supports that ezGeno can facilitate users to train genomic models and help realize the mechanism of gene regulation more.

## Notes

https://github.com/ailabstw/ezGeno.

### Competing Interest Statement

The authors have declared no competing interest.

